# Transcriptome Analysis of *xa5*-mediated Resistance to Bacterial Leaf Streak in Rice (*Oryza sativa* L.)

**DOI:** 10.1101/727115

**Authors:** Xiaofang Xie, Zhiwei Chen, Huazhong Guan, Yan Zheng, Jing Zhang, Mingyue Qin, Weiren Wu

## Abstract

Bacterial leaf steak (BLS) caused by *Xanthomonas oryzae* pv. oryzicola (*Xoc*) is a devastating disease in rice production. The resistance to BLS in rice is a quantitatively inherited trait, of which the molecular mechanism is still unclear. It has been proved that *xa5*, a recessive bacterial blast resistance gene, is the most possible candidate gene of the QTL *qBlsr5a* for BLS resistance. To study the molecular mechanism of *xa5* function in BLS resistance, we created transgenic lines with RNAi of *Xa5* (LOC_Os05g01710) and used RNA-seq to analyze the transcriptomes of a *Xa5*-RNAi line and the wild-type line at 9 h after inoculation with *Xoc*, with the mock inoculation with water as control. The results showed that *Xa5*-RNAi could (1) increase the resistance to BLS as expected from *xa5;* (2) alter (mainly up-regulate) the expression of hundreds of genes, most of which were related to disease resistance; and (3) greatly enhance the response of thousands of genes to *Xoc* infection, especially of the genes involved in cell death pathways, suggesting that *xa5* displays BLS resistance effect probably mainly because of the enhanced response of the cell death-related genes to *Xoc* infection.

## Introduction

Bacterial leaf streak (BLS) is a disease caused by the gram-negative bacterial pathogen *Xanthomonas oryzae* pv. oryzicola (*Xoc*) in rice. BLS is one of the most devastating quarantine diseases (Niño-Liu *et al*. 2006) in the main rice-producing areas of the world and can cause significant yield loss. BLS resistance is quantitatively inherited in rice (Tang *et al*. 2000). In contrast to qualitative disease resistance, which is controlled by single resistance (R) genes, race-specific and easily defeated by co-evolving pathogens (Kou and Wang 2010), the quantitative disease resistance is driven by multiple genes, generally non-race-specific and much more durable than qualitative resistance (Pink 2002; Wisser *et al*. 2005; Kliebenstein and Rowe 2009; Poland *et al*. 2009). Therefore, breeding of disease-resistant cultivars is an effective strategy to control BLS. However, such effort has often been compromised due to lack of understanding of the underlying molecular mechanism for BLS resistance.

To date, at least 13 quantitative trait loci (QTLs) conferring BLS resistance have been mapped in rice (Tang *et al*. 2000; Zheng *et al*. 2005; Chen *et al*. 2006). In addition, a recessive gene *bls1* showing race-specific resistance to BLS (Wen-Ai *et al*. 2012) and a locus *Xo1* conferring complete resistance to the African clade of *Xoc* strains of BLS (Triplett *et al*. 2016) are also reported. Interestingly, a non-host resistance gene *Rxo1* from maize also displays qualitative resistance to BLS in rice (Zhao *et al*. 2005), which specifically activates multiple defense pathways related to hypersensitive response (HR) against *Xoc*, including some signaling pathways and basal defensive pathways such as the ethylene (ET) and salicylic acid (SA) pathways (Zhou *et al*. 2010).

Overexpression of a resistance protein differentially expressed protein gene 1 (DEPG1) that contains a nucleotide-binding site-leucine rich repeat (NBS-LRR) domain results in increased susceptibility to *Xoc* strain RS105 and inhibition of some genes related to basal defensive pathways, implying its role of negative regulation for immunity in rice (Guo *et al*. 2012). Similarly, suppression of some defense related (DR) genes, such as *OsWRKY45-1* (Tao *et al*. 2009), *OsMPK6* (Shen *et al*. 2010) and *NRRB* (Guo *et al*. 2014), also presents negative regulation for immunity, increasing resistance to BLS. On the contrary, overexpression of genes *OsPGIP4, GH3-2*, and *OsHSP18.0-CI* significantly enhances resistance to BLS in rice, implying their positive regulation roles for resistance to *Xoc* (Fu *et al*. 2011; Feng *et al*. 2016; Ju *et al*. 2017).

In our laboratory, a major QTL *qBlsr5a* conferring BLS resistance was previously mapped on rice chromosome 5 (Tang *et al*. 2000) and further fine mapped to a 30-kb interval (Xie *et al*. 2014). Three genes were annotated in the interval. Among them, LOC_Os05g01710 was considered to be the most possible candidate gene, which encodes a transcription initiation factor IIA gamma (TFIIAγ) protein. There were two nucleotides different between the LOC_Os05g01710 allele from the resistant parent and that from the susceptible parents, resulting in a substitution at the 39^th^ amino acid between their encoding proteins (Xie *et al*. 2014). Interestingly, the allele of LOC_Os05g01710 from the resistant parent was identical to *xa5* in sequence, a recessive resistance gene against bacterial blight caused by *Xanthomonas oryzae* pv. oryzae (Xoo) (Iyer and McCouch 2004), implying that suppression of the dominant allele of LOC_Os05g01710 (*Xa5*) from the susceptible parent would increase the BLS resistance if it is really the gene responsible for the *qBlsr5a* effect. This implication has been verified by the result of RNA interference (RNAi) of *Xa5* (Yuan *et al*. 2016), in which the *Xa5*-RNAi lines display enhanced resistance to BLS. Therefore, it is rational to infer that *xa5* is the cause of the BLS-resistance effect of *qBlsr5a*, although the possibility of contribution from the other two genes in the interval cannot be completely excluded.

The present study aimed to investigate the molecular mechanism of *xa5*-mediated resistance to BLS in rice by analyzing the effects of *Xa5*-RNAi on gene expression regulation and transcriptional response to BLS pathogen *Xoc*. We revealed that *Xa5*-RNAi could greatly alter (mainly up-regulate) the expression of many genes related to disease resistance and enhance transcriptional response to *Xoc*, and *Xa5* affects BLS resistance probably mainly by regulating the expression of genes involved in cell death.

## Materials and methods

### Plant materials

An *indica* rice cultivar Minhui 86 (MH86) and a *japonica* rice cultivar Nipponbare were used in the experiment. Both cultivars were susceptible to BLS.

### Vector construction and rice transformation

To construct *Xa5*-RNAi (abbreviated as XR) vector, a 515-bp fragment containing the whole coding sequence of LOC_Os05g01710 (*Xa5*) was amplified from the cDNA of Nipponbare by PCR. The amplified fragment was inserted into the pTCK303 vector downstream of the ubiquitin promoter, in both forward and reversed orientations (Figure 1). The vector was introduced into *Agrobacterium* strain EHA105 using the freeze-thaw method, and further into the calli of Nipponbare and MH86 derived from mature embryos following the protocol of (Nishimura *et al*. 2006). Positive transgenic lines were identified by PCR using the hygromycin specific primers. Ten positive T2 plants from each line were selected for qRT-PCR analysis and BLS-resistance assessment. All the primers used for XR vector construction and transgenic line identification are shown in Table 1.

**Figure 1.**
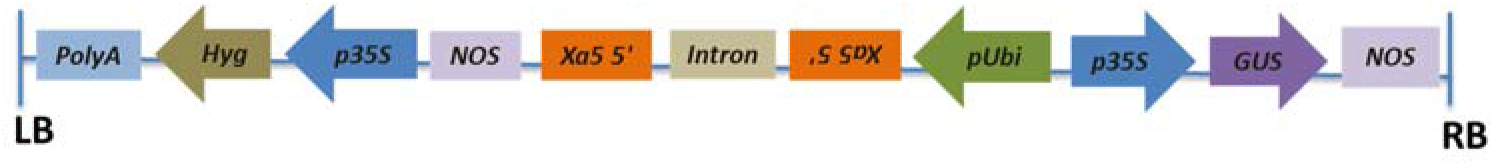
Schematic diagram of *Xa5*-RNAi construct

**Table 1.**
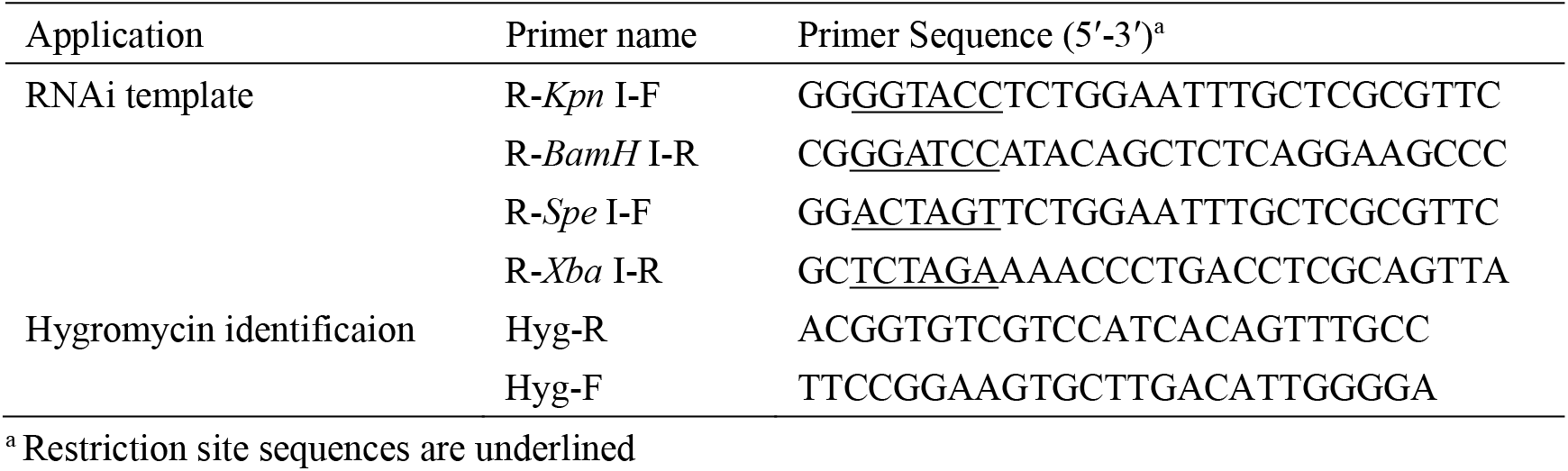
Primers used in vector construction and transgenic plant identification

### Pathogen infection and resistance assessment

The *Xoc* strain used for inoculation was kindly provided by Prof. Guoying Cheng of Huazhong Agricultural University. Rice plants were inoculated using the pricking inoculation method (Tang *et al*. 2000) at the active tillering stage and lesion length was scored from three leaves of each plant 15 days after inoculation. The resistance ability of a line was indicated by the mean lesion length of 10 plants.

### Quantitative real-time PCR analysis

Total RNA of leaves was extracted using TRIzol reagent (Invitrogen, http://www.invitrogen.com). First-strand cDNA synthesis was performed using PrimeScript™ RT reagent kit with gDNA eraser (Takara, Japan) following the manufacturer’s instruction. The cDNA samples were then assayed by qRT-PCR using SYBR Premix Ex Taq (Takara). The gene-specific primers used for qRT-PCR analysis are listed in Table S1. *Actin* was used as an internal control. Two or more biological replicates and three technical replicates were tested. The relative expression level of a gene was calculated using the 2^-ΔΔCt^ method (Livak and Schmittgen 2001). The paired t-test method was used to examine the difference of gene expression level between different samples.

### RNA sequencing and data analysis

Leaves of wild-type (WT) MH86 and one of its XR lines (M-4) at the active tillering stage were inoculated with *Xoc* or sterile water (mock inoculation, as control) and total RNA was extracted from the leaves 9 h after inoculation. In total, there were four treatments: MH86 mock (denoted as WTM), MH86 inoculated (WTI), M-4 mock (XRM), and M-4 inoculated (XRI). Three biological replicates were set for each treatment. Therefore, there were 12 RNA samples in total. Each RNA sample was from a mixture of three 1-cm leaf segments next to the inoculation site. The RNA samples were sequenced on Illumina HiSeq™ 2000 performed by Biomarker Technologies (http://www.biomarker.com.cn/, BioMarker, Beijing, China). Reads (100 bp in length, paired-end) were mapped to the reference genome and genes available at the Rice Genome Annotation Project (http://rice.plantbiology.msu.edu). For gene expression analysis, the numbers of matched reads were normalized by the RPKM (reads per kb per million mapped reads) method (Mortazavi *et al*. 2008). DEGs were identified using the criteria of |log_2_(fold change)| ≥ 2 and false discovery rate (FDR) ≤ 0.01. Gene ontology (GO) analysis was performed based on the Gene Ontology Database (http://www.geneontology.org/). The MapMan tool (http://MapMan.gabipd.org) was used for a graphical overview of pathways involving the DEGs.

## Results

### XR plants and their BLS resistance

In total, 13 and 9 independent transgenic (T_0_) XR plants were obtained from Nipponbare and MH86, respectively, and 8 T_2_ XR lines were subsequently derived, with 4 from Nipponbare (N-1/2/3/4) and MH86 (M-1/2/3/4) each. qRT-PCR analysis indicated that *Xa5* expression was significantly reduced in the XR lines (Figure 2c). Inoculation test showed that the lesion length in the XR lines was also significantly decreased in comparison with that in WT plants (Figure 2a and b). The lesion length and the *Xa5* expression level were significantly correlated, with a correlation coefficient of 0.93 (P < 0.05) in the Nipponbare XR lines and 0.94 (P < 0.05) in the MH86 XR lines, respectively. These results reconfirmed that *xa5* can enhance BLS resistance, and suggested again that *xa5* is the cause of BLS-resistance of QTL *qBlsr5a*.

**Figure 2.**
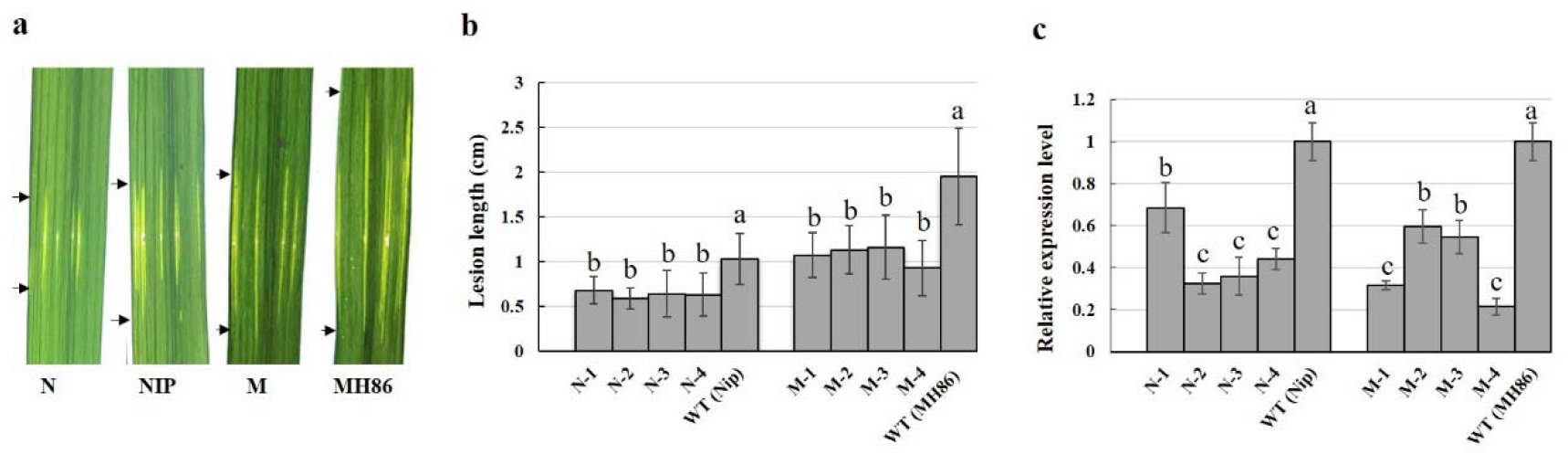
Performance of BLS resistance in *Xa5*-RNAi lines from Nipponbare and MH86. (a) Leaves of WT and RNAi lines with BLS lesions at 15 d after *Xoc* inoculation. (b) Lesion lengths of WT and T2 RNAi lines measured at 15 d after *Xoc* inoculation. (c) Relative expression levels of *Xa5* in WT and T2 RNAi lines at 15 d after *Xoc* inoculation. RNAi lines of N-1/2/3/4 and M-1/2/3/4 were derived from Nipponbare and MH86, respectively. Letters indicate statistical significance of the difference at 0.01 level according to ANOVA-Tukey’s test.

### Reads of RNA sequencing

Since the XR line M-4 from MH86 showed the most significant suppression of *Xa5* expression and increase of BLS resistance (Figure 2), it was used for transcriptome analysis by RNA sequencing (RNA-seq) together with MH86. In total, 12 RNA samples were sequenced. More than 50 million reads were obtained in each sample, of which most (>80%) were uniquely mapped to the Nipponbare reference genome (Table 2). These uniquely mapped reads were used for subsequent gene expression analysis. The RNA-seq data have been submitted to the database of the NCBI Sequence Read Archive (http://trace.ncbi.nlm.nih.gov/Traces/sra) under the accession number PRJNA558068.

**Table 2.** Statistics of RNA-seq reads

| Samples | Total reads | Uniquely mapped reads | Multiply mapped reads |
| --- | --- | --- | --- |
| XRI-1 | 59,395,708 | 50,385,642 (84.83%) | 1,834,255 (3.09%) |
| XRI-2 | 71,452,986 | 60,635,913 (84.86%) | 1,366,235 (1.91%) |
| XRI-3 | 61,645,792 | 49,929,674 (80.99%) | 1,141,129 (1.85%) |
| XRM-1 | 54,530,318 | 47,518,520 (87.14%) | 922,469 (1.69%) |
| XRM-2 | 59,370,030 | 52,035,796 (87.65%) | 1,343,567 (2.26%) |
| XRM-3 | 62,654,504 | 52,749,774 (84.19%) | 1,346,979 (2.15%) |
| WTI-1 | 52,052,314 | 45,482,023 (87.38%) | 1,326,379 (2.55%) |
| WTI-2 | 60,142,368 | 52,009,381 (86.48%) | 1,723,060 (2.86%) |
| WTI-3 | 58,057,590 | 50,885,173 (87.65%) | 1,148,390 (1.98%) |
| WTM-1 | 57,207,506 | 49,886,919 (87.20%) | 1,121,745 (1.96%) |
| WTM-2 | 51,845,292 | 45,544,161 (87.85%) | 1,018,135 (1.96%) |
| WTM-3 | 53,018,932 | 46,897,761 (88.45%) | 891,019 (1.68%) |

Note: Reads were mapped to the Nipponbare reference genome.

To validate the RNA-seq data, qRT-PCR (Table S1) was used to examine the expression of 6 genes encoding proteins related to disease resistance (including phytohormones, phyto-oxygenase, etc.) in the four treatments. The expression patterns of these six genes detected by qRT-PCR were consistent with those detected by RNA-seq (Figure 3), indicating that the RNA-seq data were reliable.

**Figure 3.**
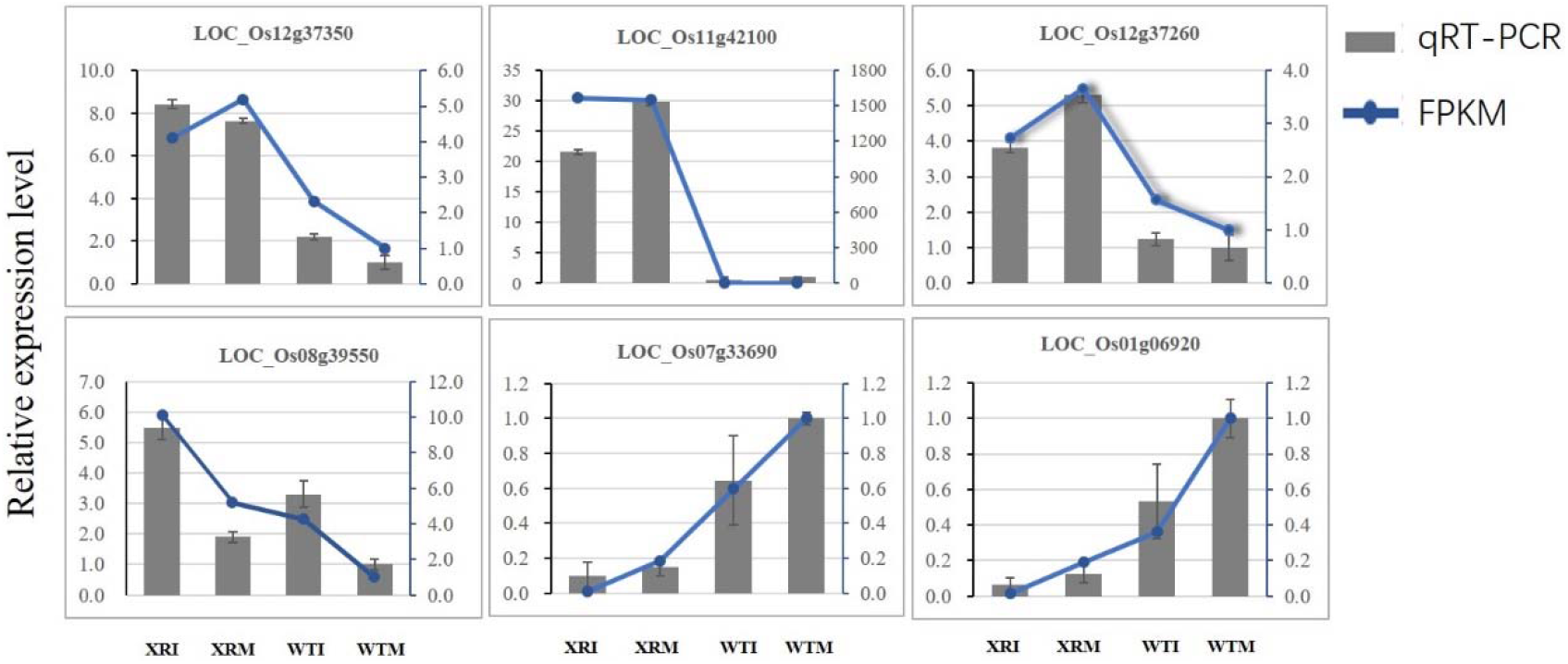
Quantitative RT-PCR analysis of six genes to validate the RNA-seq data.

### Differentially expressed genes

We analyzed differentially expressed genes (DEGs) in three comparisons, namely, C1: XRM vs. WTM; C2: WTI vs. WTM; and C3: XRI vs. XRM. C1 was used to examine the effect of XR on gene expression, while C2 and C3 were used to examine the effect of XR on transcriptional response to *Xoc* infection.

A total of 317 DEGs were detected in C1 (Table S2), among which ~3/4 (244) genes were up-regulated, suggesting that *Xa5* functions mainly as a negative regulator for many genes. MapMan analysis showed that many of the up-regulated genes were involved in the biological pathways related to biotic stress (Figure S1), including hormone signaling, cell wall, beta glucanase, proteolysis, defense genes (DR), redox state, signaling, Myb, secondary metabolites, etc. Most of these genes, such as those associated with cell wall function, phytohormone of salicylic acid (SA), jasmonic acid (JA) and ethylene (ET), have been found to play important roles in plant defenses against pathogens (Yang *et al*. 2015), including *Xoc* in rice (Shen *et al*. 2010; Guo *et al*. 2014; Feng *et al*. 2016).

Figure S1 MapMan overview of biotic stress showing the transcriptional changes in C1 (XRM vs. WTM). Individual genes are represented by small squares, where red and blue indicate significant up-regulation and down-regulation, respectively. The color scale displays log2-transformed fold changes.

There were 157 DEGs (125 up-regulated + 32 down-regulated) in C2 (Table S3) and 3,115 DEGs (1,202 up-regulated + 1,913 down-regulated) in C3 (Table S4), respectively. The number of DEGs in C3 was ~19 times more than that in C2, suggesting that XR can dramatically enhance the transcriptional response to *Xoc* infection. However, the set of DEGs in C3 only covered ~1/3 (56/157) of that in C2, although most of overlapped DEGs showed the same expression change directions in C2 and C3 (Figure 4). In addition, while there were more up-regulated genes than down-regulated genes in WT (C2), the trend was reversed in XR (C3; Figure 4). These results suggested that XR can also significantly alter the pattern of transcriptional response to *Xoc* infection.

**Figure 4.**
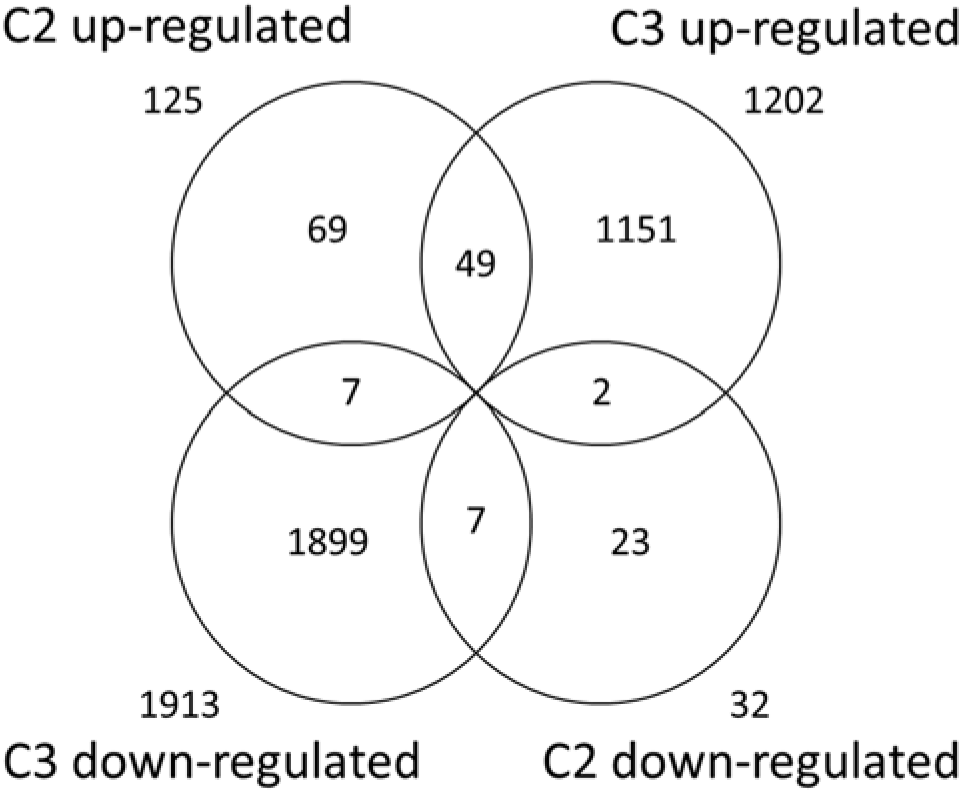
Venn diagram of DEGs detected by RNA-seq in C2 (WTI vs. WTM) and C3 (XRI vs. XRM).

### Gene ontology enrichment

Gene ontology (GO) analysis showed that the DEGs in C1 were significantly enriched (*P*-value < 0.01) in 43 GO terms (Table S5), including 29 terms on biological process (BP), 1 on cellular component (CC) and 13 on molecular function (MF). Among the BP terms, ~3/4 (22/29) were known to be related to plant disease resistance, which could be classified into several groups, including oxidation-reduction (Matika and Loake 2014), siderophore (Aznar and Dellagi 2015), secondary metabolite (Piasecka *et al*. 2015), cell death (Coll *et al*. 2011) and so on. This was consistent with the result of MapMan analysis, suggesting that a main role of *Xa5* is to regulate the expression of genes related to disease resistance.

In C2, 78 GO terms were significantly enriched (*P*-value < 0.01) with DEGs (Table S6), including 41 on BP, 9 on CC and 28 on MF, respectively. Almost all of the BP terms were related to disease resistance, which could be also classified into several groups similar to those observed in C1 except for the group of cell death.

The DEGs in C3 were enriched (*P*-value < 0.01) in 66 GO terms (Table S7), including 18 terms on BP, 21 on CC and 27 on MF, respectively. Among the BP terms, eight were related to plant disease resistance, which could be classified into several groups, including cell death (3 terms), defense response (1 term), thiamine metabolism (4 terms) and so on. Thiamine is known to function as an activator of plant disease resistance (Il-Pyung *et al*. 2005). The cell death group terms were the most significant, with *P*-values (≤ 1.1×10^-11^) at least 10^5^ times smaller than any other term, implying that this group is particularly important for the function of *Xa5* on BLS resistance.

Surprisingly, although there were much more DEGs in C3 than in C2, the number of enriched GO terms on BP in C3 was much smaller than that in C2. Comparison showed that there were no common BP terms between C2 and C3. However, they had 16 and 4 BP terms common with C1, respectively, all of which were related to disease resistance (Figure 5).

**Figure 5.**
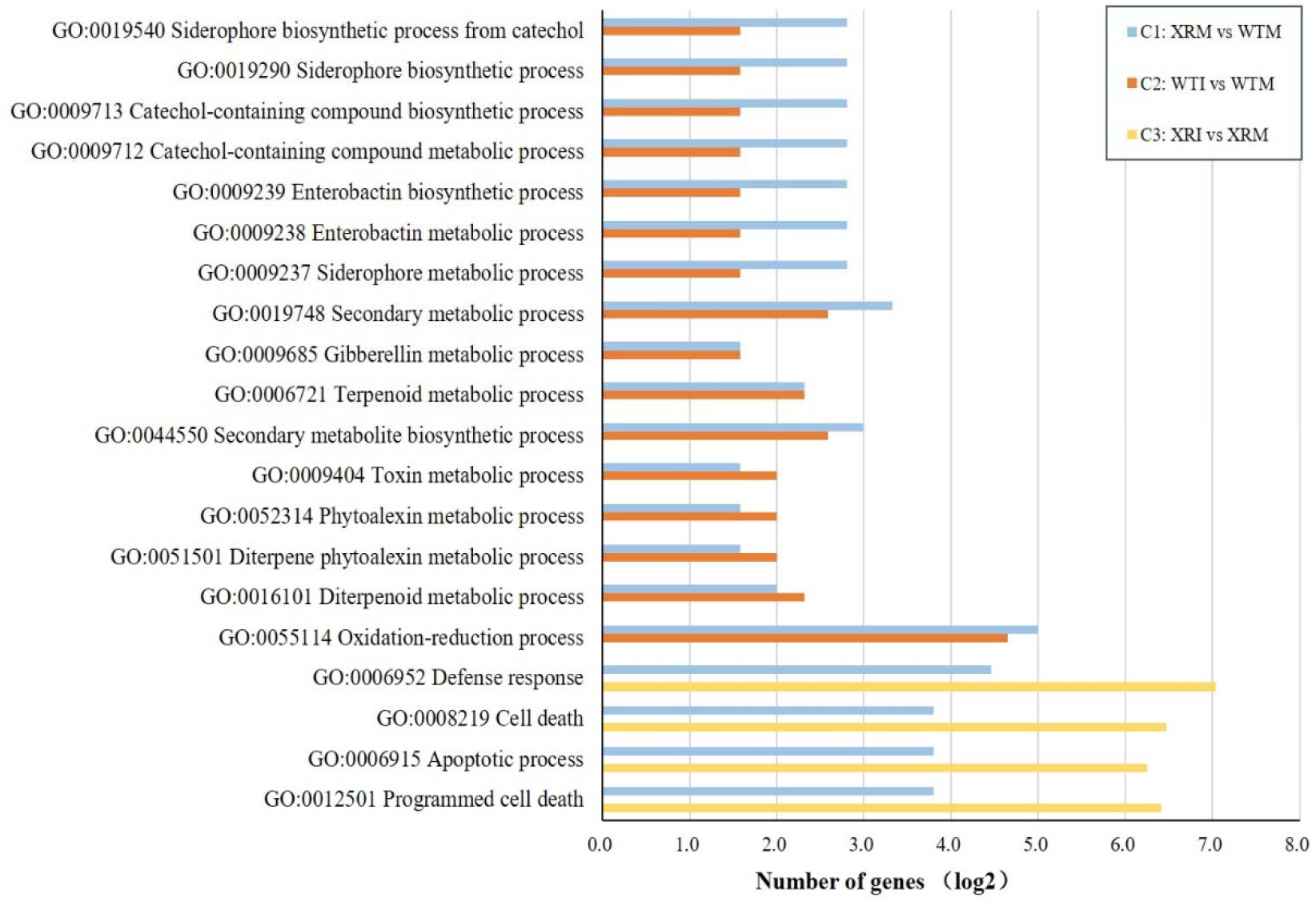
Significant GO terms on biological process common in C1, C2 and C3.

## Discussion

TFIIAγ is a component of the general transcription factor IIA and is involved in all polymerase II-dependent transcription in eukaryotes (Li *et al*. 1999; Høiby *et al*. 2007). Two TFIIAγ-like genes, TFIIAγ1 and TFIIAγ5 (*Xa5*), have been found in rice. The recessive allele *xa5*, which encodes a V39E substitution variant of TFIIAγ5, is functionally confirmed to enhance the resistance to rice bacterial blight (Iyer and McCouch 2004; Jiang *et al*. 2006). In addition, the results of fine mapping of QTL *qBlsr5a* (Xie *et al*. 2014) and RNAi of *Xa5* (Yuan *et al*. 2016) have suggested that *xa5* is the most possible candidate gene of the QTL *qBlsr5a* conferring resistance to BLS. This study verified the *Xa5*-RNAi effect on BLS resistance (Figure 2), further affirming the inference that *Xa5* is responsible for the effect of *qBlsr5a* on BLS resistance.

It is generally considered that plants have evolved two different innate immune systems: the pathogen-associated molecular pattern (PAMP)-triggered immunity (PTI) system and the effector-triggered immunity (ETI) system (Jones and Dangl 2006). PTI belongs to a relatively weak immune response triggered by PAMPs, which depends on the basal defense to restrict the colonization of invading pathogens (Zipfel *et al*. 2014). As PTI has no race specificity, it is predicted to confer durable and broad spectrum resistance (Liu *et al*. 2014). ETI is mediated by polymorphic resistance proteins that can recognize the highly variable effectors from pathogens, with a fast and strong response usually accompanied with a hypersensitivity reaction (HR) and eventually programmed cell death (PCD) to restrict biotrophic cellular pathogens (Dodds and Rathjen 2010). We have seen above that the DEGs in C1 were mainly enriched in four groups of GO terms on BP related to disease resistance, namely, oxidation-reduction, siderophore, secondary metabolite, and cell death (Table S5). Among these GO terms, the former three groups belong to the mechanisms of basal defense, while the last group (cell death) belongs to the mechanism of HR. Therefore, they are responsible for PTI and ETI, respectively. This suggests that *Xa5* regulates both PTI- and ETI-related genes.

It is interesting that some of the GO terms related to disease resistance in C1 were also detected in the enrichment analyses in C2 and C3, but the PTI-related terms only detected in C2, while the ETI-related terms only detected in C3, and there was no overlap between C2 and C3 (Figure 5). In addition, the ETI-related terms were very highly significant and also the most significant in C3 (Table S7). These results suggest that it is likely that the *xa5*-mediated BLS resistance is mainly due to the response of the genes involved in the cell death pathways to *Xoc* infection. It appears that the dominant allele *Xa5* can inhibit the response of these genes to *Xoc* infection. Therefore, no GO terms about cell death could be detected in C2.

Certainly, PTI-related genes may also contribute to disease resistance. But why the above-mentioned PTI-related GO terms were not significant in C3? The possible reason could be that the response of the genes in these PTI-related GO terms to *Xoc* infection is similar to that to *Xa5*-RNAi (Figure 5), and the effects of these two types of response are not additive but superimposed. Thus, since the response of the genes to *Xa5*-RNAi has already existed in an XR line, the response to *Xoc* infection is masked and therefore becomes undetectable.

In conclusion, as a component of a general transcriptional factor, *Xa5* plays an important role in the regulation of many genes related to disease resistance (including both PTI-related and ETI-related genes) in rice. Suppression of its expression can lead to defense-oriented reprogramming and thereby limit the multiplication or spread of the BLS pathogen *Xoc* through a stronger and more direct immune response like ETI to protect the plant. This defense strategy could accompany with a similar occurrence of a hypersensitivity reaction (HR), and eventually resulted in programmed cell death (PCD), cell death, or apoptotic to against the pathogen invasion.

## Authors’ contributions

Conceived and designed the experiments: WW. Performed the experiments: XX ZC HG YZ JZ MQ. Analyzed the data: XX. Contributed materials: ZC. Wrote the paper: XX WW. All authors read and approved the manuscript.

